# Arf6 anchors Cdr2 nodes at the cell cortex to control cell size at division

**DOI:** 10.1101/2021.08.17.456680

**Authors:** Hannah E. Opalko, Kristi E. Miller, Hyun-Soo Kim, Cesar Vargas-Garcia, Abhyudai Singh, Michael-Christopher Keogh, James B. Moseley

**Author notes:** Correspondence to James B. Moseley.

## Abstract

Fission yeast cells prevent mitotic entry until a threshold cell surface area is reached. The protein kinase Cdr2 contributes to this size control system by forming multiprotein nodes that inhibit Wee1 at the medial cell cortex. Cdr2 node anchoring at the cell cortex is not fully understood. Through a genomic screen, we identified the conserved GTPase Arf6 as a component of Cdr2 signaling. Cells lacking Arf6 failed to divide at a threshold surface area and instead shifted to volume-based divisions at increased overall size. Arf6 stably localized to Cdr2 nodes in its GTP-bound but not GDP-bound state, and its GEF (guanine nucleotide exchange factor) Syt22 was required for both Arf6 node localization and proper size at division. In *arf6Δ* mutants, Cdr2 nodes detached from the membrane and exhibited increased dynamics. These defects were enhanced when *arf6Δ* was combined with other node mutants. Our work identifies a regulated anchor for Cdr2 nodes that is required for cells to sense surface area.

## INTRODUCTION

Many cell types maintain constant size during recurring cycles of growth and division. Such size control indicates the existence of mechanisms that delay cell cycle transitions until cells reach a critical size threshold (Amodeo and Skotheim, 2016; Rupes, 2002). The fission yeast *Schizosaccharomyces pombe* is a strong model system to study size-dependent control of entry into mitosis, also called the G2/M transition. These rod-shaped cells grow by linear extension and enter mitosis and divide at a reproducible size (Wood and Nurse, 2015). Although cell length, volume, and surface area scale together during linear extension growth, recent studies show that *S. pombe* cells divide at a specific surface area, as opposed to length or volume (Pan et al., 2014; Facchetti et al., 2019). Thus, the underlying size control network likely operates through mechanisms connected to the cell cortex.

The G2/M transition is controlled by regulated activation of cyclin-dependent kinase Cdk1 (Harashima et al., 2013). Multiple mechanisms contribute to cell size-dependent Cdk1 activation in *S. pombe*. The protein kinase Wee1 phosphorylates and inhibits Cdk1 in small cells (Russell and Nurse, 1987a; Gould and Nurse, 1989), while a regulatory network progressively inhibits Wee1 as cell size increases (Lucena et al., 2017; Allard et al., 2018; Opalko et al., 2019). The concentrations of mitotic inducers including Cdc13 (mitotic cyclin) and Cdc25 (protein phosphatase that counteracts Wee1) increase as cells grow (Moreno et al., 1990; Keifenheim et al., 2017; Patterson et al., 2019). The Wee1 regulatory network draws particular interest because it functions at the cell cortex through the conserved protein kinases Cdr1 and Cdr2. Mutations in *cdr1* and *cdr2* cause cells to divide at a larger size due to uninhibited Wee1 (Russell and Nurse, 1987b; Young and Fantes, 1987; Breeding et al., 1998; Kanoh and Russell, 1998). In addition to increased cell size, *cdr2Δ* mutants fail to divide according to surface area and instead shift towards volume-based size control, meaning that Cdr1/2-Wee1 signaling links cell surface area with the G2/M transition (Facchetti et al., 2019).

Cdr1 and Cdr2, members of the conserved SAD-family (Synapses of amphids defective) protein kinases, play distinct roles in Wee1 inhibition. Cdr1 directly phosphorylates the Wee1 kinase domain to inhibit its catalytic activity (Coleman et al., 1993; Parker et al., 1993; Wu and Russell, 1993; Opalko et al., 2019). Cdr2 does not regulate Wee1 kinase activity directly and instead provides spatial control to this pathway. Cdr2 forms oligomeric “nodes,” which are stable structures tethered to the plasma membrane and positioned in the cell middle (Morrell et al., 2004). Cdr2 recruits both Cdr1 and Wee1 to these sites, and Cdr2 kinase activity correlates with the dwell time of Wee1 at individual nodes (Moseley et al., 2009; Martin and Berthelot-Grosjean, 2009; Allard et al., 2018). Since Cdr2 is progressively activated as cells grow, Wee1 spends more time at nodes as cells grow larger, resulting in its cell size-dependent phosphorylation and inhibition (Deng et al., 2014; Allard et al., 2018). Importantly, this mechanism acts at the plasma membrane, consistent with its role connecting cell surface area with the G2/M transition. Cdr2 nodes also function after mitotic entry by serving as spatial landmarks for cytokinetic ring assembly (Almonacid et al., 2009).

Nodes are critical to Cdr1/2-Wee1 signaling, so it is important to identify mechanisms of node formation and anchoring at the cell cortex. Cdr2 contains a KA1 domain that is required for node formation in cells and binds to lipids, in part through an RKRKR motif (Rincon et al., 2014; Morrell et al., 2004). The Cdr2 KA1 domain also displays some clustering activity, while Cdr2-interacting factors, including anillin-like Mid1, contribute to Cdr2 clustering in a KA1-independent manner (Rincon et al., 2014). These node-forming activities are countered by the protein kinase Pom1, which directly phosphorylates Cdr2 to inhibit interactions with the membrane and Mid1 (Rincon et al., 2014). Pom1 is concentrated at cell tips to restrict Cdr2 node assembly to the medial cell cortex (Bähler and Pringle, 1998; Moseley et al., 2009; Martin and Berthelot-Grosjean, 2009). Understanding how Cdr2 oligomerizes into nodes and interacts with the plasma membrane is crucial to explaining how Wee1 is regulated by cell surface area.

In this study we identify the conserved GTPase Arf6 as a novel component and regulator of Cdr1/2-Wee1 nodes. Loss of Arf6 impairs signaling to Wee1 and leads to Cdr2 clusters that are detached from the plasma membrane. Genetic experiments indicate that Arf6 anchors nodes at the cortex in parallel to other mechanisms including the Cdr2 KA1 domain and Cdr2-Mid1 interactions. The role of Arf6 in assembling these multicomponent signaling platforms at the plasma membrane has implications for kinase clusters in other systems.

## RESULTS AND DISCUSSION

### Arf6 regulates the cell cycle through the Cdr2 pathway

The Cdr2 pathway inhibits Wee1, and *cdr1Δ* and *cdr2Δ* mutants are synthetically lethal in combination with the temperature-sensitive *cdc25-22* mutation (Russell and Nurse, 1987b; Young and Fantes, 1987; Breeding et al., 1998; Kanoh and Russell, 1998). To screen for new components of Cdr2 signaling (Fig 1A), we individually combined a library of 3,004 viable *S. pombe* deletion mutants (~75% of the non-essential fission yeast genome) with *cdc25-22* using synthetic genetic array analysis (SGA; as in (Roguev et al., 2007)) (Fig 1B-C). As controls, we combined each library deletion with either a *cdr2Δ* mutant or a *cdc25+* wild type allele. We reasoned that mutations in Cdr2 signaling should exhibit synthetic sick / synthetic lethal (SS/SL) interactions with *cdc25-22*, but should be non-additive with *cdr2Δ*, while *cdc25+* controlled for linkage effects. From the resulting double mutants, 66 were SS/SL with only *cdc25-22*, 31 mutants were SS/SL with only *cdr2Δ*, and 28 were SS/SL with both *cdc25-22* and *cdr2Δ* (Fig 1D, Table S1).

**Fig 1:**
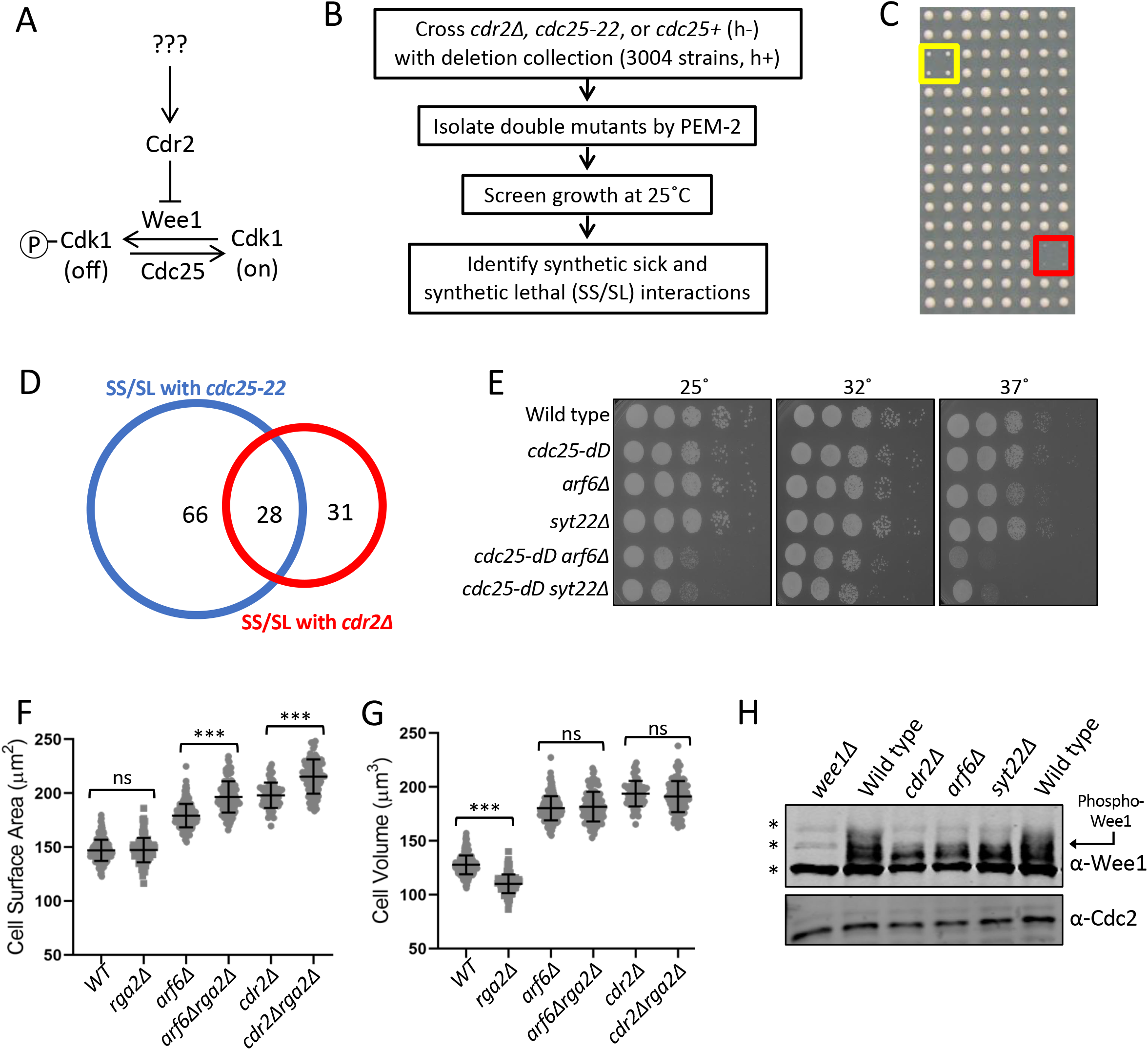
Arf6 regulates cell size through the Cdr2-Wee1 pathway. (A) Schematic of the Cdr2-Wee1 pathway. (B) SGA workflow (see Methods). (C) Example of SGA growth screening. All strains were spotted in quadruplicate. Yellow box shows slow growth; red box shows no growth. (D) Venn diagram of SGA hits. (E) Serial dilution assay of the indicated strains. (F) Surface area of dividing cells from the indicated strains. (G) Volume of dividing cells from the indicated strains. n ≥ 70 cells per strain; ns p ≥ 0.05; *** p ≤ 0.001. Graphs show mean ± SD. (H) Western blot of whole cell extracts for endogenous Wee1. Asterisks indicate background bands; Cdc2 is loading control.

Mutants in the Cdr2 pathway exhibit altered cell length at division, and this phenotype is synthetic with mutations in *cdc25* (Breeding et al., 1998; Kanoh and Russell, 1998). Thus, we were interested to find that *arf6Δ* was SS/SL with *cdc25-22* and increased cell length at division when combined with *cdc25-22* (Fig S1A). We decided to focus on Arf6, a conserved GTPase linked in animal cells to formation of multivalent protein assemblies at the plasma membrane (Cherfils, 2014), the site of Cdr2 nodes in fission yeast. In both serial dilution growth assays and analysis of cell length at division, *arf6Δ* was non-additive with *cdr2Δ* but showed synthetic defects with the hypomorphic allele *cdc25-dD* (Figs 1E, S1B-D). GTPase signaling is activated by GEFs (guanine nucleotide exchange factors) and inhibited by GAPs (GTPase-activating proteins). Deletion of the Arf6 GEF (*syt22Δ*) phenocopied *arf6Δ* genetic interactions with *cdr2Δ* and *cdc25-dD* (Figs S1B-D). Previous studies identified a role for fission yeast Arf6 and its regulatory GEF (Syt22) and GAP (Ucp3) in bipolar cell growth (Fujita, 2008; Fujita and Misumi, 2009, 2011), but a role in cell cycle progression has not been reported. Our data suggest a novel function for active Arf6 in the Cdr2 cell cycle pathway.

The cell length at division phenotype for *arf6Δ* was minor, but these cells were wider than wild type (Fig S1E). Due to this increased width, the enlarged cell size phenotype of *arf6Δ* was nearly as severe as *cdr2Δ* when comparing cell surface area (Fig 1F). Recent work using cell width mutants showed that fission yeast cells divide at a specific surface area, as opposed to length or volume (Facchetti et al., 2019; Pan et al., 2014). In addition, *cdr2Δ* mutants shift to dividing at a set volume that is larger than wild type cells (Facchetti et al., 2019). By comparing wild type and *rga2Δ*, which has reduced cell width (Villar-Tajadura et al., 2008), we confirmed that fission yeast cells divide at a specific surface area, not volume (Fig 1F-G). Perhaps more importantly, *arf6Δ* cells and *arf6Δ rga2Δ* cells divided at the same volume but at distinct surface areas, similar to *cdr2Δ* and *cdr2Δ rga2Δ* (Fig 1F-G). Thus, *arf6Δ* cells (like *cdr2Δ*) divide at a larger size and shift from surface area to volume-based divisions.

The molecular output of the Cdr2 pathway is inhibitory phosphorylation of Wee1, which can be monitored by SDS-PAGE mobility shifts (Allard et al., 2018, 2019; Opalko et al., 2019). The slower migrating, hyperphosphorylated form of Wee1 was lost in *arf6Δ* and *syt22Δ*, similar to *cdr2Δ* (Fig 1H). We conclude that activated Arf6 functions in the Cdr2 pathway to control cell size at division through inhibition of Wee1.

### Regulated localization of Arf6 to nodes

Next, we tested if Arf6 functions at nodes by examining its localization. Arf6-mNeonGreen (mNG) localized to the plasma membrane as previously shown (Fujita, 2008), but was strongly enriched at cortical nodes in the cell middle (Fig 2A). Arf6 and Cdr2 were highly colocalized at nodes (Fig 2B, S2A), identifying Arf6 as a new node component. The Arf6-mNG signal at individual nodes was stable by time lapse microscopy (Fig 2C), similar to Cdr2 and Cdr1 but distinct from Wee1 (Pan et al., 2014; Allard et al., 2018). To test the timing of Arf6 node localization, we tracked mitosis with the spindle pole body marker Sad1 (Hagan and Yanagida, 1995), and tracked cytokinesis with the myosin-II regulatory light chain Rlc1 (Naqvi et al., 2000; Le Goff et al., 2000). Arf6 localized to nodes throughout interphase but left nodes after cells entered mitosis (Fig 2D), similar to Cdr2 (Morrell et al., 2004). Arf6 returned to the cell middle during cytokinesis but did not constrict with the ring. Following cell separation, Arf6 reappeared at nodes. Arf6 node localization required Cdr2 but not other node proteins (Fig 2E, S2B). These data show that Arf6 is a component of Cdr2 nodes in addition to functioning in the Cdr2 pathway.

**Fig 2:**
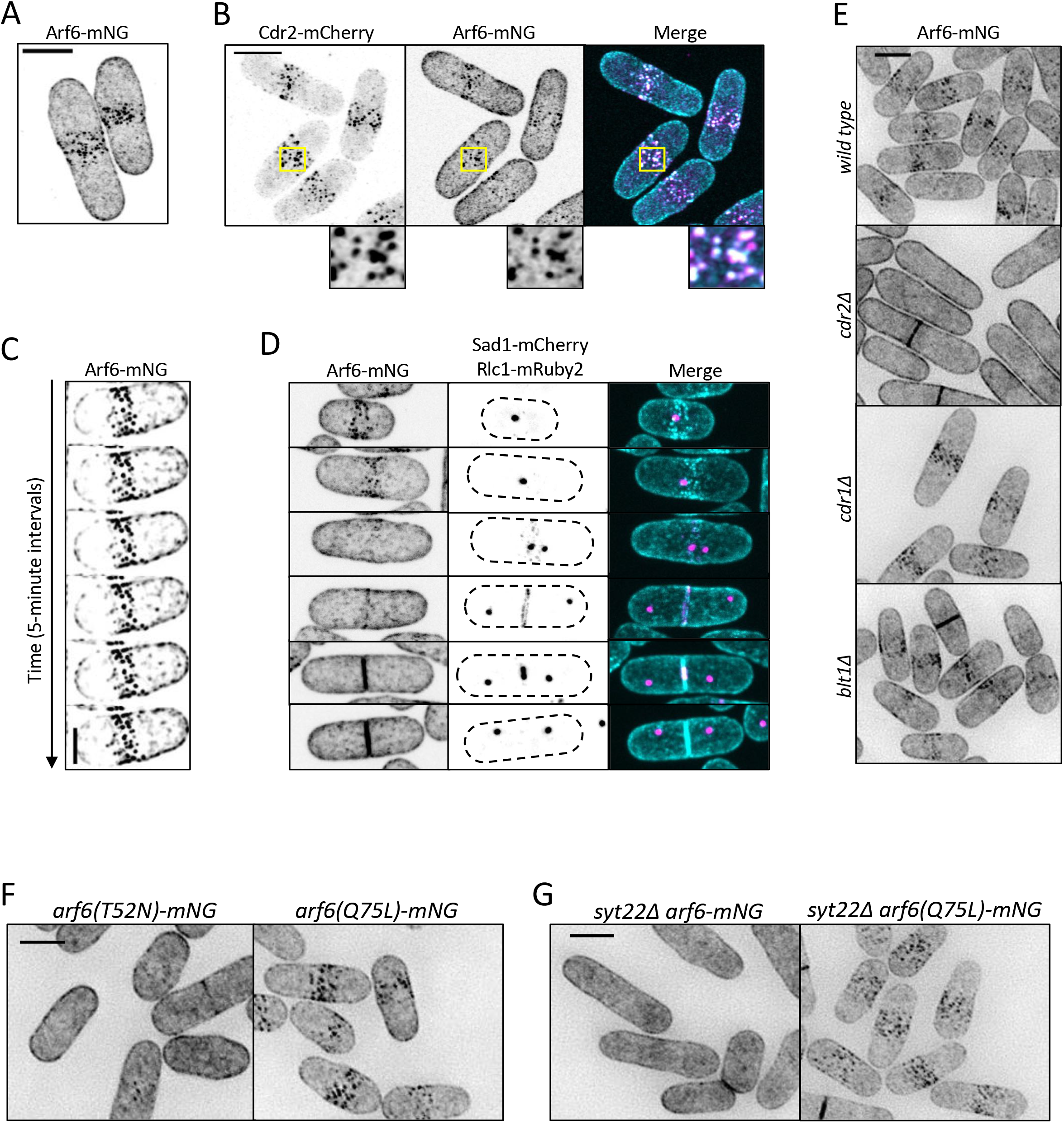
Arf6 is a novel node component. (A-B) Maximum intensity projection of Airyscan images. Insets show zoom of boxed area. (C) Maximum intensity projections of deconvolved z series from images taken every 5 minutes. Scale bar is 3 μm. (D) Localization of Arf6 at different cell cycle stages. Sad1 and Rlc1 mark mitosis and cytokinesis respectively. Maximum intensity projections of z series. (E) Localization of Arf6-mNG in the indicated mutants. Images are sum intensity projections of deconvolved z series. (F-G) Sum intensity projections of deconvolved z series. Scale bars in A-B and D-G are 5 μm.

We next asked how GTP-binding and hydrolysis regulate Arf6 localization to nodes. A GDP-locked mutant arf6(T52N)-mNG lost node localization, but the GTP-locked allele arf6(Q75L)-mNG remained at nodes (Fig 2F). Further, Arf6 localization to nodes was lost upon deletion of its GEF Syt22 (Fig 2G). The Arf6 GAP Ucp3 is essential (Fujita and Misumi, 2011), so we could not test its role in Arf6 localization. If the Arf6 localization defect in *syt22Δ* is due to loss of the GTP-bound state, then it should be suppressed by *arf6(Q75L)*. Indeed, arf6(Q75L)-mNG localized to nodes even in *syt22Δ* cells (Fig 2G). These data show that GTP binding is required for Arf6 localization to nodes and suggest that Arf6 localizes to nodes in its GTP-bound state. Loss of Arf6 from Cdr2 nodes explains *syt22Δ* defects in cell size and Wee1 phosphorylation.

Next, we tested if Arf6 localized to nodes with its GEF (Syt22) or GAP (Ucp3). Syt22-mNG localized to cortical puncta as previously shown (Fig S2C) (Fujita and Misumi, 2009). Syt22 puncta were spread throughout the cell periphery and did not colocalize with Cdr2 (Fig S2C-D). Ucp3-mNG localized to spots at the cell tips, which likely represent endocytic actin patches due to colocalization with actin patch component Pan1 (Fig S2E-F). We conclude that Syt22 and Ucp3 do not localize to nodes with Arf6, but they regulate Arf6 node localization through its nucleotide-bound state.

Arf6 GTPases have a conserved myristoylated glycine and an alpha helix that work together to promote membrane binding (Gillingham and Munro, 2007) (Fig S2G). To test if membrane binding contributes to Arf6 node localization, we made a non-myristoylated *arf6* mutant (G2A) and two stepwise helix deletions (N3-S10Δ and K11-F17Δ) because complete helix deletion abrogated expression. The resulting *arf6* alleles reduced node localization, and instead enriched at the cytoplasm (Fig S2H-I). Therefore, reducing Arf6 membrane binding also reduces Arf6 node localization.

Together, these results show that Arf6 localizes stably to Cdr2 nodes during interphase in a manner that depends on nucleotide binding, membrane binding, and Cdr2 itself.

### Arf6 promotes cortical tethering of Cdr2 nodes

How does Arf6 promote Cdr2 node function? We imaged Cdr2-mEGFP in *arf6Δ* cells and observed several node defects. First, *arf6Δ* cells had cytoplasmic clusters of Cdr2 that were absent in wild type cells (Fig 3A, S2J), indicating defects in cortical anchoring. Second, Cdr2 was present at cell tips in addition to nodes in *arf6Δ* cells (Fig S2K-L). Third, Cdr2 node intensity was reduced in *arf6Δ* cells (Fig S2M). The brightest nodes, which are lacking in *arf6Δ* cells, are likely diffraction-limited clusters of smaller/unitary nodes (Akamatsu et al., 2017). Fourth, timelapse imaging revealed reduced stability of Cdr2 signal at some but not all nodes in *arf6Δ* cells (Fig 3B, S3A). In addition, FRAP experiments showed increased Cdr2 dynamics at nodes of *arf6Δ* cells consistent with loss of anchoring (Fig 3C, S3B), but slower Cdr2 dynamics at nodes in the GTP-locked mutant *arf6(Q75L)* consistent with hyper-stable anchoring (Fig S3C). These defects indicate that Arf6 anchors Cdr2 stably at nodes, meaning that Arf6 and Cdr2 reciprocally promote each other’s node localization.

**Fig 3:**
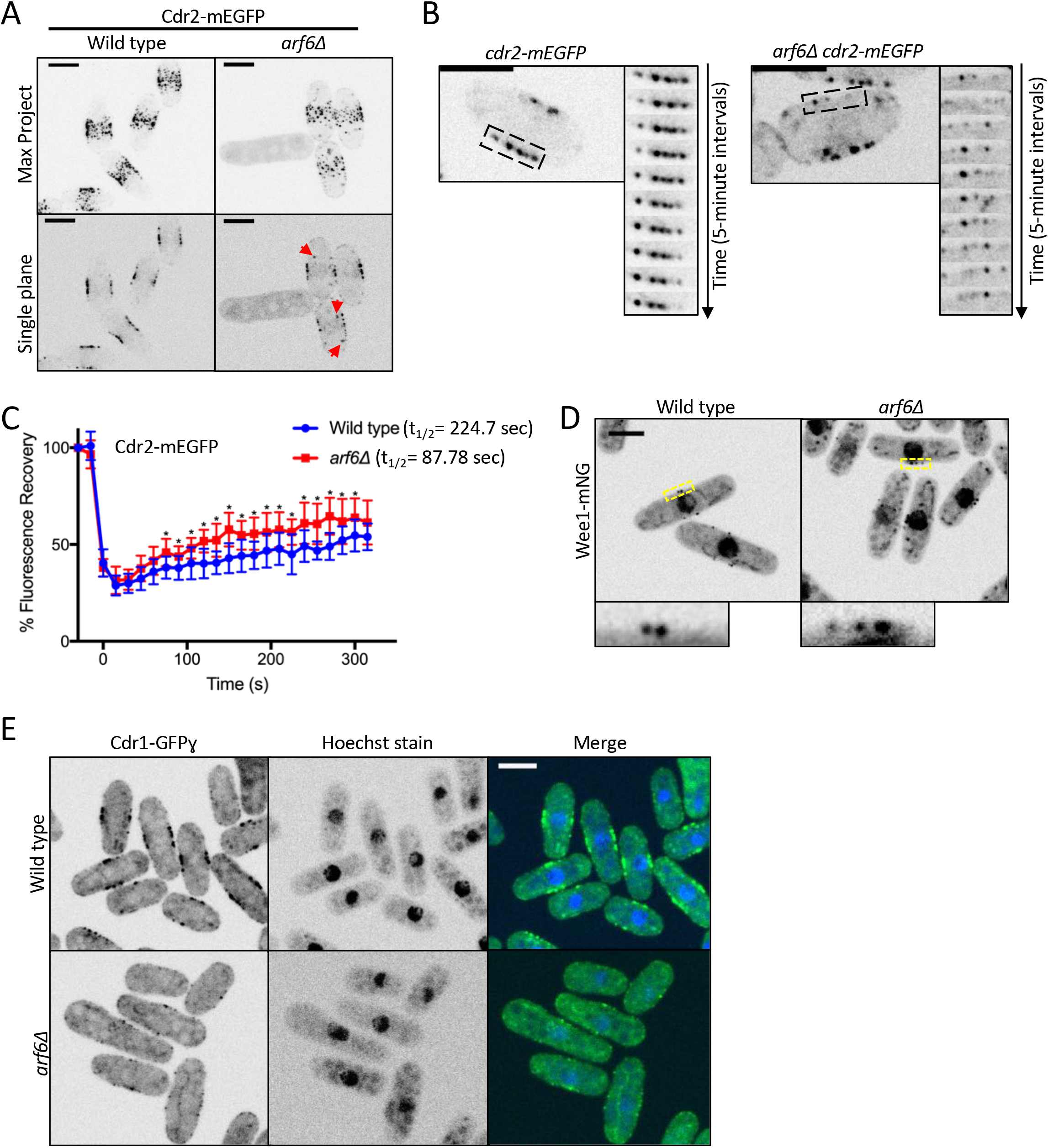
Cdr2 nodes are disrupted in *arf6*Δ. (A) Maximum intensity projections of z series (top) or single middle focal planes (bottom). Red arrows indicate cytoplasmic Cdr2 puncta. (B) Time lapse imaging of Cdr2-mEGFP from middle focal plane. Montages show 5-minute timepoints from boxed regions. (C) FRAP analysis of Cdr2-mEGFP in wildtype and *arf6Δ* cells. n = 10 cells each. Points are mean ± SD, * p ≤ 0.05. (D) Middle-focal plane images of Wee1-mNG. Insets are zoom of yellow boxes. (E) Middle focal plane images. All scale bars 5 μm.

Node defects in *arf6Δ* correlate with loss of Wee1 phosphorylation and altered cell size. To explain this connection, we examined the localization of Wee1 and Cdr1 at cortical nodes in *arf6Δ* mutants. Wee1 localized to nodes in *arf6Δ* but Cdr1 did not (Figs 3D-E, S3D). Loss of Cdr1 from Cdr2 nodes was most obvious in line traces along the cortex of *arf6Δ cdr1-3xGFP cdr2-RFP* cells (Fig S3E-F). Because the 3xGFP tag on Cdr1 reduces cell size, we also tested Cdr1 localization in *arf6Δ* cells elongated with the *cdc25-dD* mutation and observed similar defects (Fig 3E). We conclude that loss of node integrity in *arf6Δ* prevents recruitment of Cdr1, leading to loss of Wee1 inhibitory phosphorylation and cell size defects.

We next tested how Arf6 connects with three other mechanisms that contribute to Cdr2 node anchoring. First, Cdr2 has a membrane-binding KA1 domain required for cortical localization. Mutation of an RKRKR motif in the KA1 domain reduced Cdr2 node localization (Rincon et al., 2014), and this defect was enhanced by *arf6Δ* (Fig S3G). Second, the node protein Mid1 interacts with Cdr2 to promote clustering into nodes (Rincon et al., 2014). We combined *arf6Δ* with the *mid1(400-450Δ)* mutant that cannot bind Cdr2. In the resulting cells, Cdr2 was absent from the cell cortex and formed large cytoplasmic puncta (Fig 4A, S2J). Thus, Arf6 and Mid1 are partially overlapping anchors for Cdr2 nodes. We note that the Cdr2 KA1 domain was still present in these cells but was not sufficient for cortical localization. Third, we combined *arf6Δ* with mild overexpression of Pom1, which phosphorylates Cdr2 to reduce its membrane binding, catalytic activity, and clustering (Deng et al., 2014; Bhatia et al., 2014; Rincon et al., 2014). *P3nmt1-pom1* alone causes mild defects in Cdr2 localization (Moseley et al., 2009), but *arf6Δ P3nmt1-pom1* largely eliminated Cdr2 from the cortex and caused localization to cytoplasmic puncta (Fig 4B, S2J). We conclude that Arf6 contributes to Cdr2 cortical anchoring together with parallel mechanisms, most notably Mid1 and Pom1.

**Fig 4:**
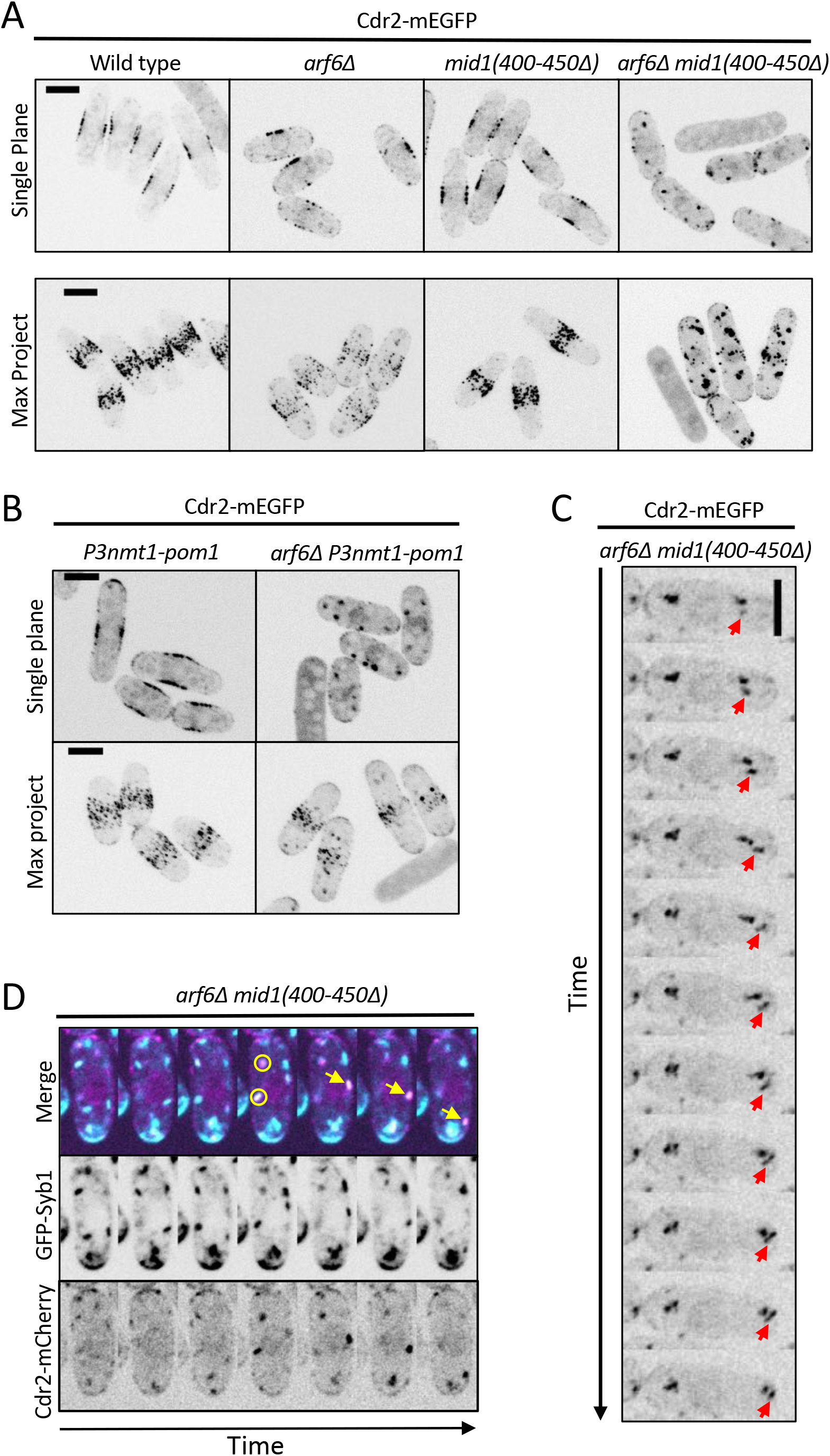
Cdr2 node mutants exacerbate *arf6Δ* phenotypes. (A-B) Cdr2-mEGFP localization in the indicated strains. Single focal planes and maximum intensity projections images of z series. (C) Montage of a nodelay time lapse for single focal plane. Arrow follows a moving Cdr2 punctum. (D) Mobile Cdr2 puncta colocalize with the exocytic vesicle marker Syb1. Circles mark colocalizing puncta, arrow marks moving colocalized punctum. Cells were imaged every 30 seconds. All scale bars 5 μm.

By time lapse microscopy, we observed motile cytoplasmic Cdr2 puncta in 14 out of 20 *arf6Δ mid1*(*400-450Δ*) cells analyzed, with most movement directed towards cell tips (Fig 4C). These Cdr2 puncta colocalized with Syb1, the v-SNARE synaptobrevin that marks exocytic vesicles, as puncta moved towards cell tips in *arf6Δ mid1*(*400-450Δ*) (Fig 4D). Cdr2 did not colocalize with Syb1 in wild type cells (Fig S3H). These data suggest that Cdr2 becomes enriched on exocytic vesicles when it cannot be anchored at cortical nodes by Arf6 and Mid1.

### Arf6 contributes to cytokinesis

Cdr2 contributes to cytokinesis by recruiting anillin-like protein Mid1 to nodes during interphase (Almonacid et al., 2009). The Mid1 N-terminus (Mid1-Nter) is necessary and sufficient for its cytokinesis function, and the localization and function of Mid1-Nter requires Cdr2 (Celton-Morizur et al., 2006; Almonacid et al., 2009). Unlike the Mid1(Δ400-450) mutant that cannot interact with Cdr2, the Mid1-Nter mutant interacts with Cdr2 and depends on Cdr2 nodes for cytokinesis, meaning that Mid1-Nter provides a system to examine the function of Cdr2 nodes in cytokinesis. Given the Cdr2 node anchoring defects in *arf6Δ*, we tested the localization and function of Mid1-Nter in *arf6Δ*. In *arf6+* cells, GFP-Mid1-Nter localized to the cortex to support proper cell morphology and division plane positioning (Fig 5A). In contrast, GFP-Mid1-Nter mislocalized in *arf6Δ* cells, which were multi-nucleated with grossly aberrant septa (Fig 5A, S3I-J). In addition, Cdr2-mEGFP did not concentrate in medial nodes in *arf6Δ mid1*-Nter cells (Fig 5B, S3K). These data indicate that Arf6 is required for the localization and function of Mid1-Nter in cytokinesis.

**Fig 5:**
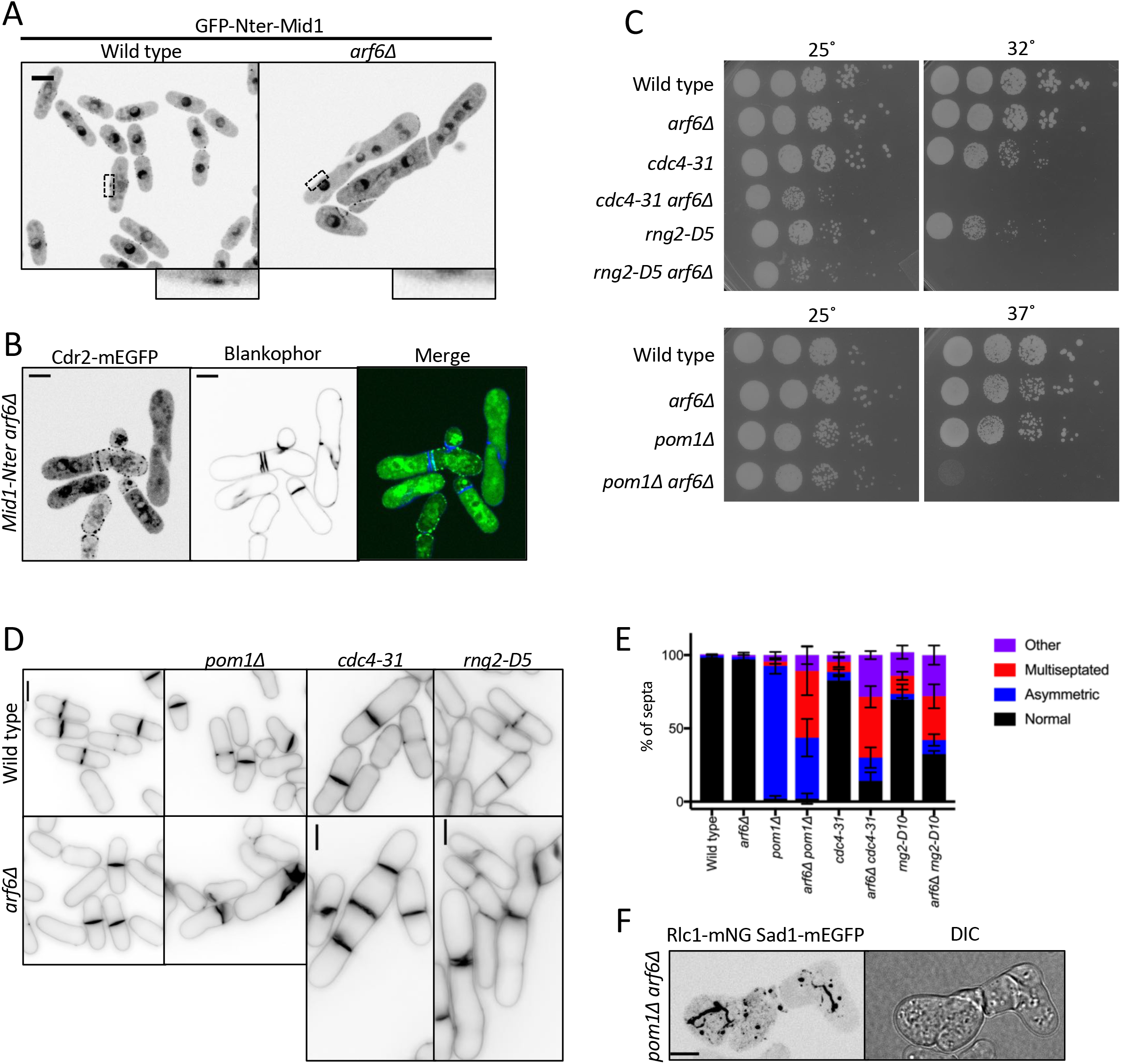
Arf6 promotes robust cytokinesis. (A) Middle focal plane images of GFP-Nter-Mid1. Insets are zoom of black boxes. (B) Cdr2-mEGFP localization in *mid1-Nter arf6Δ* cells stained with blankophor. (C) Serial dilution growth assays for the indicated strains. (D) Blankophor images of the indicated strains. (E) Quantification of septation defects in the indicated strains. Values are means ± SD from 3 biological replicate experiments. (F) Localization of Rlc1 and Sad1 in representative *arf6Δ pom1Δ* cells. Maximum intensity projection of a z series. All scale bars 5 μm.

We wondered if this result reflected a broad role for Arf6 in promoting the robustness of cytokinesis. We did not observe defects in the timing of cytokinetic ring formation, maturation, or constriction in *arf6Δ* cells (Fig S3L), so we tested genetic interactions between *arf6Δ* and several cytokinesis mutants. We combined *arf6Δ* with *cdc4-31*, a temperature-sensitive mutant in the myosin-II essential light chain (McCollum et al., 1995); with *rng2-D5*, a temperature-sensitive mutant in IQGAP (Eng et al., 1998; Chang et al., 1996); or with *pom1Δ*, which positions the division plane asymmetrically (Bähler and Pringle, 1998). All three double mutants exhibited synthetic defects in cell growth and cytokinesis as judged by tilted and disorganized septa (Figs 5C-E, S3M). We attempted to visualize cytokinesis in *arf6Δ pom1Δ* cells using Rlc1-mNG and Sad1-mEGFP to mark the actomyosin ring and SPBs respectively (Fig 5F). However, strong disorganization of Rlc1 in these cells prevented meaningful time lapse microscopy experiments. We conclude that Arf6, like other node proteins, contributes to robust cytokinesis.

## Conclusions

Our study identifies the conserved GTPase Arf6 as a new functional component of Cdr2 nodes. Human ARF6 functions in several membrane-localized processes such as endocytosis, lipid homeostasis, and cytokinesis (Schweitzer et al., 2011; Gillingham and Munro, 2007), and has been linked to the proliferation, invasion, and metastasis of multiple cancers (Hashimoto et al., 2004; Li et al., 2017). A general theme of Arf GTPases is the assembly and stabilization of multivalent membrane-bound platforms (Cherfils, 2014). We demonstrated a clear role for Arf6 in organizing nodes for cell size control. We found that Cdr2 nodes lose stability in *arf6Δ* cells, and unstable nodes fail to recruit Cdr1 leading to reduced Wee1 phosphorylation and cell size defects. These findings suggest that Arf6 promotes stable node coalescence by promoting multivalent protein-protein and protein-lipid interactions. In the absence of this clustered node anchoring, cells lose surface area-sensing and shift to volume-based divisions. Arf6 localization to nodes depends on the state of its bound nucleotide under control of the GEF Syt22. It will be interesting to learn how cell growth and expansion control the balanced activities of both Syt22 and the Arf6 GAP Ucp3. Because *arf6Δ* mutants fail to divide according to cell surface area and instead shift to volume-based division, it seems likely that plasma membrane expansion regulates Arf6 through Syt22 and/or Ucp3 as part of a mechanism to link cell cycle progression with overall cell size.

---

**Supplemental Figure S1.**
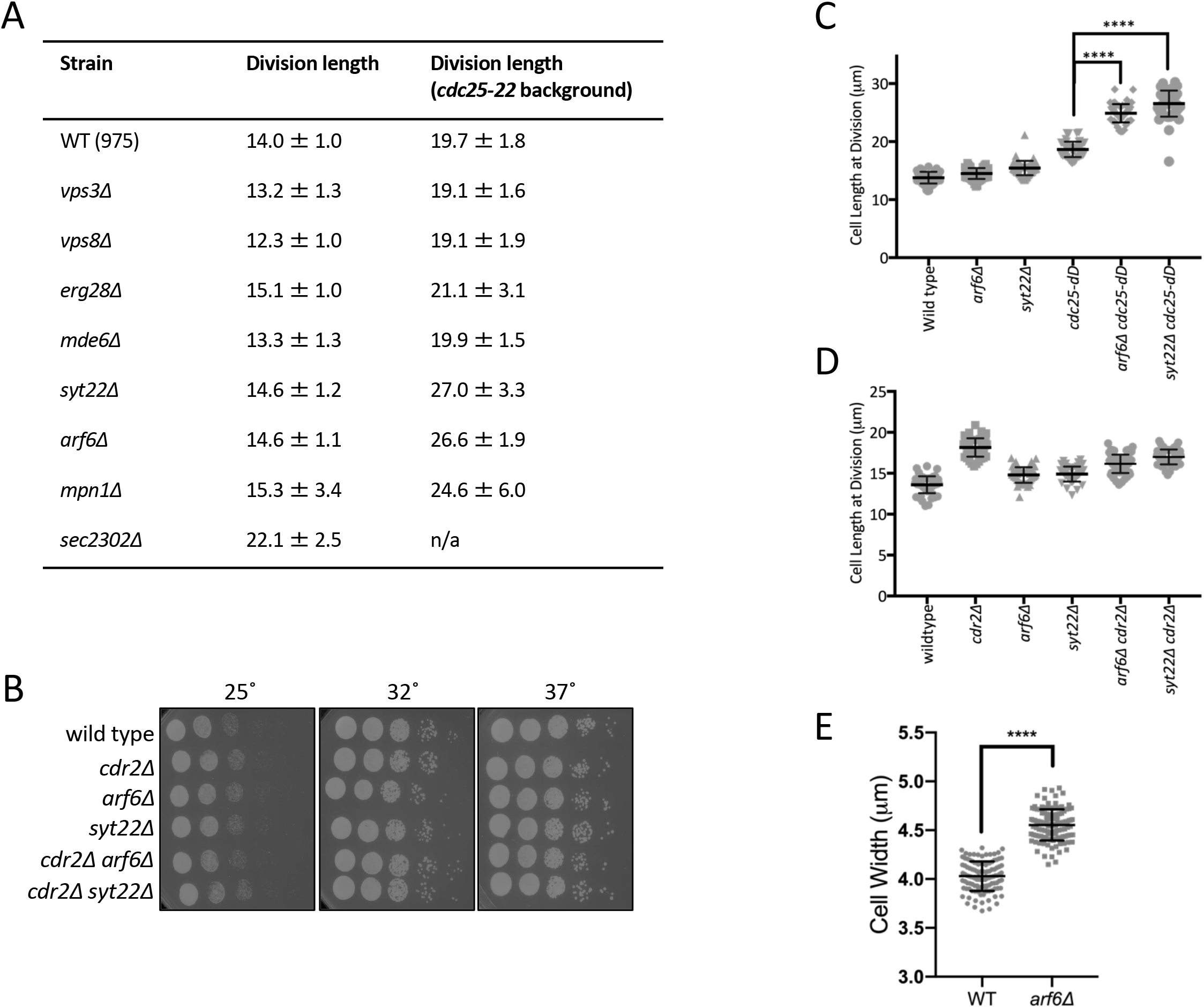
(A) Cell length at division measurements reporting mean ± SD for n>50 cells each. (B) Serial dilution growth assays. (C-D) Cell length at division for the indicated strains. n>50 cells each. Graphs show mean ± SD. (E) Cell width measurement for the indicated strains. **** p ≤ 0.0001.

**Supplemental Figure S2.**
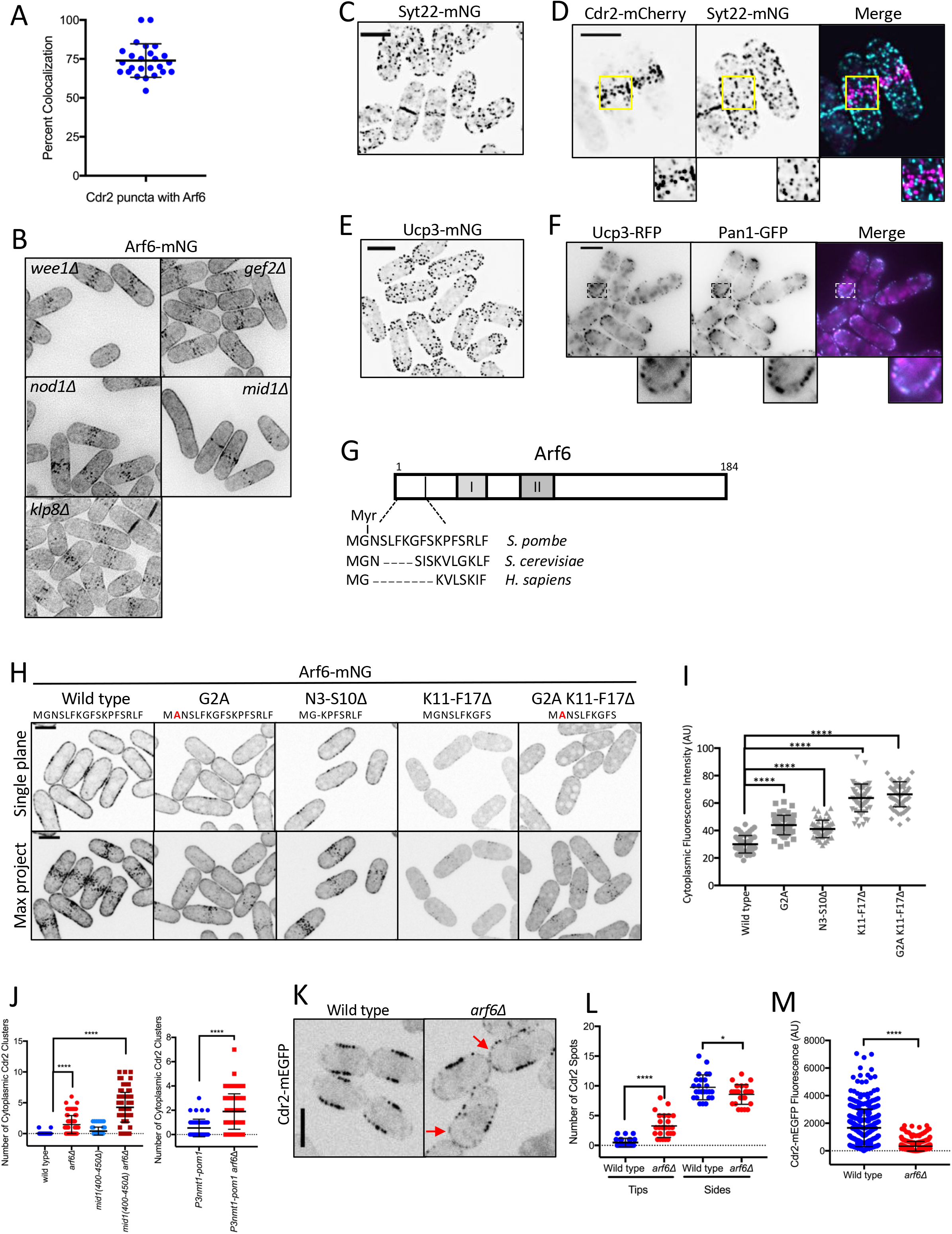
(A) Arf6 is a component of most Cdr2 nodes. Each point indicates the percentage of Cdr2 nodes with Arf6 present from a single cell. Bars represent mean ± SD. (B) Localization of Arf6 in the indicated mutants. Images are sum intensity projections of deconvolved z series. (C) Maximum intensity projection of a deconvolved z series. (D) Maximum intensity projection of a deconvolved z series. Insets show zoom of boxed area. (E) Maximum intensity projection of a deconvolved z series. (F) Middle single focal plane images. Insets show zoom of boxed area. (G) Schematic showing relative conservation of the Arf6 N-terminus from fission yeast, budding yeast, and humans. (H) Single middle focal plane and maximum intensity projections from z series. (I) Cytoplasmic fluorescence intensity of the indicated strains. n > 50 cells each. Graphs show mean ± SD. (J) Cdr2 cytoplasmic puncta counted from 5 middle focal planes. n > 50 cells each. (K) Single focal plane images. Red arrows point to Cdr2 at cell tips. (L) Quantification of Cdr2 nodes/spots at cell sides and cell tips. Note the increase in Cdr2 spots at cell tips in *arf6Δ* cells. (M) Quantification of Cdr2-mEGFP signal per node from the indicated strains. Each point represents a single node. Bars represent mean ± SD. **** p ≤ 0.0001, * p ≤ 0.05. All scale bars 5 μm.

**Supplemental figure S3.**
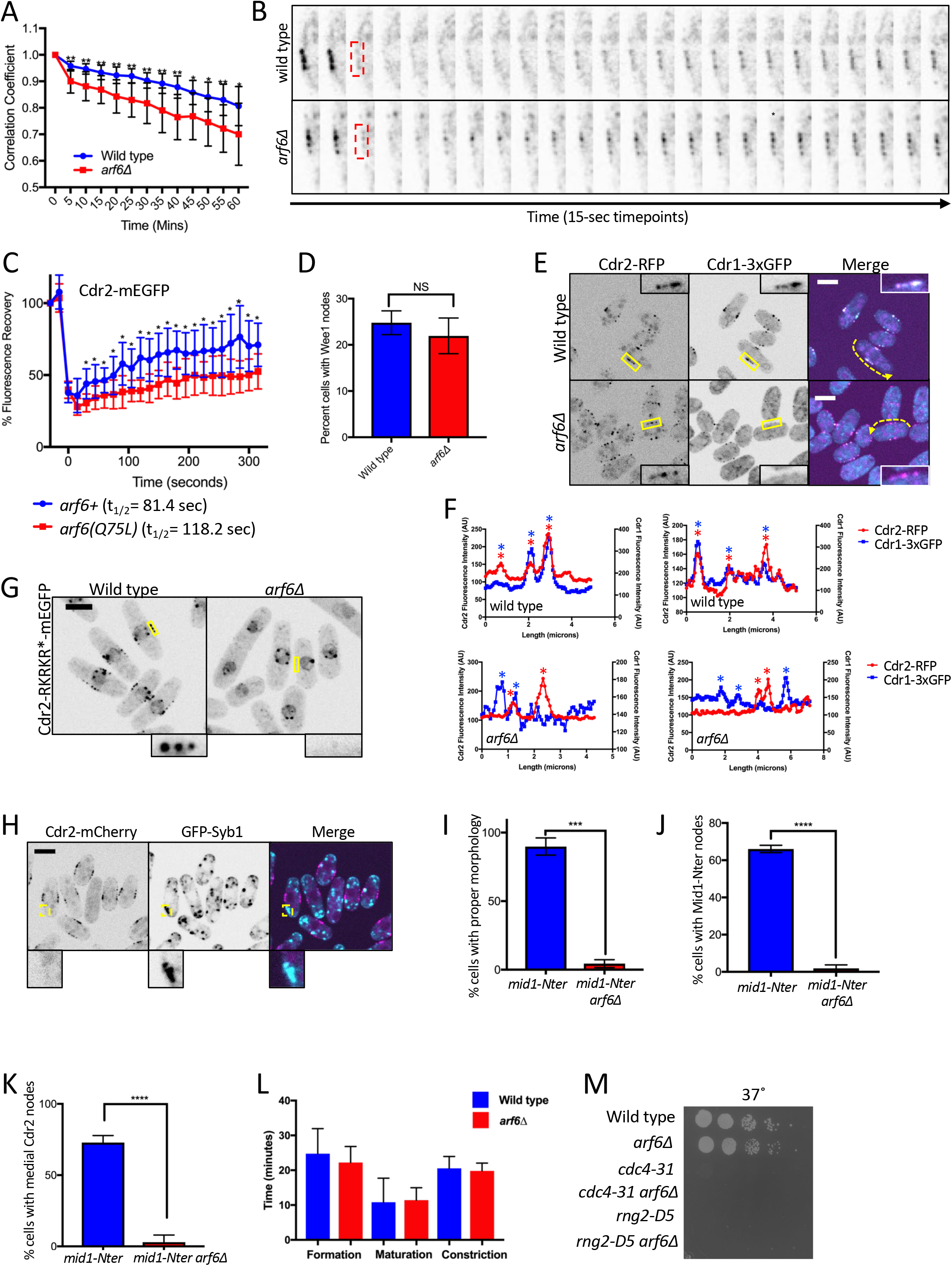
(A) Quantification of Cdr2-mEGFP dynamics from time lapse imaging. For single cells, the Pearson correlation coefficient was measured for each time point compared to the initial image (time = 0); n = 10 cells for each strain. A faster rate of decay for the correlation coefficient indicates loss of stability for Cdr2-mEGFP localization. ** p ≤ 0.01, * p ≤ 0.05. (B) Examples of FRAP experiment where red boxed region was photobleached and analyzed for recovery. (C) FRAP analysis of Cdr2-mEGFP at nodes in the indicated strains. n = 10 cells each. Points are mean ± SD, * p ≤ 0.05. (D) Percentage of cells with Wee1 localization at cortical nodes for wild type versus *arf6Δ* strains. Bars indicate mean ± SD, n=100 cells each. NS, not significant. (E) Middle focal plane images. Insets are zoom of yellow boxes. (F) Fluorescence intensity of line scans along cortex of cells as in panel E (dashed yellow lines). Asterisks mark node peaks for Cdr2 (red) and Cdr1 (blue). Note overlapping peaks in wild type but not in *arf6Δ* cells. (G) Middle focal plane images. (H) Middle focal plane images, insets are zoom of yellow boxes. (I-K) Quantification of cell morphology (I), Mid1-Nter localization (J), and Cdr2 localization (K) for the indicated strains. Bars represent mean ± SD from biological triplicate experiments with n>50 cells each. **** p ≤ 0.0001. (L) Timing of cytokinesis measured with Sad1 and Rlc1. n = 20 cells each. (M) Serial dilution growth assay at 37° (from Fig 5C). All scale bars 5 μm.

**Table S1.** List of yeast strains and SGA scores from this study.

## MATERIALS AND METHODS

### Strain construction and media

Standard *Schizosaccharomyces pombe* media and methods were used (Moreno et al., 1991). Strains and plasmids used in this study are listed in the Supplemental Table. The *cdc25-dD* mutant (*dD* abbreviates *degron-DAmP*) reduces Cdc25 levels by addition of a degradation tag (degron) in combination with truncation of the 3’UTR (Breslow et al., 2008; Deng et al., 2014). Homologous recombination was used for C-terminal tagging and gene deletions as described previously (Bähler et al., 1998). To construct GTP binding and helix mutants of Arf6, the sequence pArf6-Arf6-mNG-Tadh1 was PCR amplified from genomic DNA of strain JM6125. This PCR product was ligated into pDC99 vector using KpnI/SacII sites. Each mutation was introduced using QuikChange II site-directed mutagenesis kit (Stratagene). All constructs were Sanger sequenced for verification. pDC99 vector was then linearized using NotI and transformed into *arf6Δ::natR leu1-32 ura4-D18* (JM6565). Colonies were then selected for growth on EMM-Uri media and no growth on EMM-Leu media. For serial dilution assays, cells were spotted in 10-fold dilutions on YE4S media and grown at the indicated temperature for 3-5 days.

### Synthetic genetic array screening

We used the PEM-2 approach to combine *cdc25-22* (strain JM942), *cdr2Δ* (strain JM945), and *cdc25+* (strain JM943) with a collection of 3,004 fission yeast gene deletions (~75% of the nonessential *S. pombe* genome, noting that this collection did not include *syt22Δ*) (Kim et al., 2010). For *cdc25-22* and *cdc25+*, the natR cassette was integrated 507bp downstream of the *cdc25* stop codon. Mating, haploid selection, data acquisition, and analysis were performed as previously described (Roguev et al., 2007, 2008). Colony size was scored between 0 (no growth) and 3 (wild type growth). For each deletion mutant, synthetic growth defects with *cdc25-22* were assessed by subtracting the double-mutant colony size score when combined with *cdc25-22* from that with *cdc25+*. Similar analysis was done for *cdr2Δ*. Mutants were considered synthetic sick/lethal (SS/SL) if the synthetic growth defect was −2 or −3. All scores are provided in the Supplemental Table.

### Microscopy for cell geometry measurements

Fission yeast cells were grown at 25°C in YE4S medium to logarithmic phase for imaging. Cells were collected at 4000 rpm for 15 s, placed on a coverglass-bottom dish (P35G-1.5-20C, MatTek), and then covered with a piece of prewarmed (to 25°C) YE4S agar. Images were collected using a spinning disk confocal microscope: Yokogawa CSU-WI (Nikon Software) equipped with a 60x 1.4 NA CFI60 Apochromat Lambda S objective lens (Nikon); 405 nm, 488 nm, 561 nm laser lines; and a photometrics Prime BSI camera on an inverted microscope (Eclipse Ti2, Nikon). Multiple fields per cell type were imaged within 1 hour at room temperature and images were acquired with 27 z-stacks and 0.2 μm step.

### Cell segmentation

Brightfield images were processed for cell size analysis using a partly automated pipeline. First using ImageJ, a smoothing filter and Gaussian blur was applied to each optical section of an image to reduce image noise. We performed global thresholding by selecting a grey value cutoff to produce a binary image for each z-section. An optical section outside of the focal plane with intact boundary bands around cells was selected for further processing. The flood fill tool was used (by hand) to generate a black background so image pixel values in the background are set to 0 and cells in white are 1. Binary images were further processed by morphological erosion and subsequent dilation operators to remove white regions between cell clumps for better single cell segmentation. The paintbrush tool was used to further separate clumped cells by hand and the flood fill tool was used to delete any abnormal cells or unresolved clumps of cells. Images were also processed to remove cells along the edge of the image. The resulting binary image (the ‘cell mask’) was compared to the original brightfield image and confirmed to be an accurate representation of cell size. Next, cell masks were compared to corresponding fluorescent mtagBFP2-NLS images to identify cells containing two nuclei, indicating active division. Cells containing only a single nucleus were deleted using the flood fill tool, thus generating a binary mask of only dividing cells.

### Cell geometry measurements

For a given cell segmentation, the cell width was determined by hand measurements in ImageJ using the straight-line tool. The average cell width divided by two (cell radius) was determined for a population of cells (n>100). We assumed that each cell in a given strain has the same cell radius (average of population) for calculation of cell surface area and volume (below). In MATLAB, the cell length or cell symmetry axis of individual segmented cells (from mask of dividing cells) was identified by principal component analysis of the cloud points internal to the cell as previously described (Facchetti et al., 2019). Cell surface area or volume of individual cells was calculated in MATLAB using the equation for surface area and volume of a cylinder with hemispherical ends because of the rod-like shape of fission yeast cells. An ANOVA was performed to determine statistical differences between sets of data for cell geometry analyses.

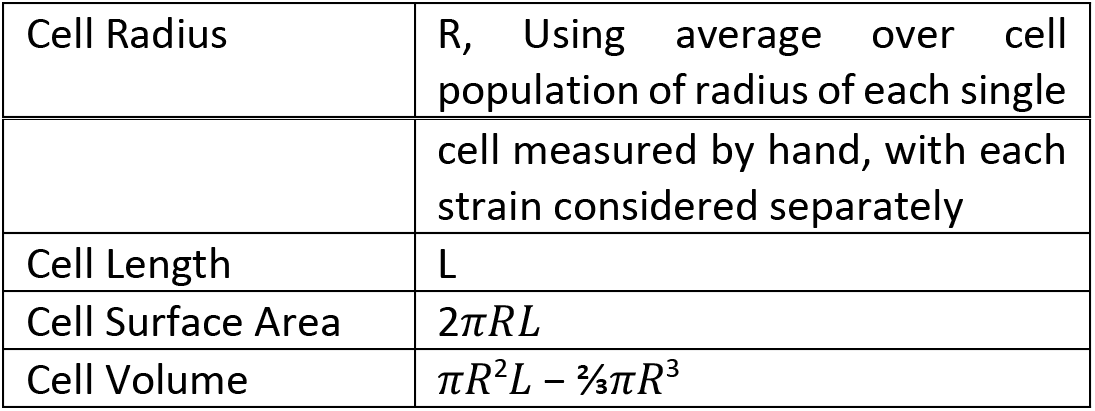

|  |  |
| --- | --- |
| Cell Radius | R, Using average over cell population of radius of each single |
|  | cell measured by hand, with each strain considered separately |
| Cell Length | $L$ |
| Cell Surface Area | $2\pi RL$ |
| Cell Volume | $\pi R^2 L - \frac{2}{3}\pi R^3$ |

### Western Blot

For western blots, 2 ODs of logarithmic phase *S. pombe* cells were flash frozen in liquid nitrogen. Whole cell extracts were made by lysing cells in 100 μl sample buffer (65mM Tris pH 6.8, 3% SDS, 10% glycerol, 10% 2-mercaptoethanol, 50 mM sodium fluoride, 50 mM β-glycerophosphate, 1 mM sodium orthovanadate, complete EDTA-free protease inhibitor tablet (Pierce)) in a Mini-beadbeater-16 for 2 minutes at 4°C, and then incubated at 99°C for 5 minutes. Lysates were briefly centrifuged to pellet insoluble material, and supernatant was isolated as whole-cell extract. Samples were resolved by reducing SDS-PAGE (10% acrylamide/bis-acrylamide for Cdc2 gels; 6% acrylamide/bis-acrylamide for all others) run at constant 20 mAmps until the 75 kDa marker approached the bottom of the gel. Gels were transferred to nitrocellulose using Trans-blot Turbo Transfer System (Bio-rad). Wee1 was probed using anti-Wee1 antibody (Allard et al., 2018). Cdc2 was used as a loading control using anti-cdc2 antibody (Santa Cruz Biotechnologies; SC-53217).

### Widefield Microscopy

For Figures 2E-G, 5D, and S2B, cells were imaged at room temperature on a DeltaVision Imaging System (Applied Precision), with an Olympus IX-71 inverted wide-field microscope, a 100x UplanSApo 1.4 NA oil objective, a Photometrics CoolSNAP HQ2 camera, and an Insight solid-state illumination unit. For images with z stacks, focal planes were imaged with 0.3 μm step size and deconvolved using SoftWoRx software (Applied Precision Ltd.) Blankophor was used to mark septating cells for cell length measurements and to highlight septum irregularities (Fig 5D). Cell length measurements, along with maximum and sum intensity projection images were created using ImageJ.

### Spinning Disc Confocal Imaging

Two spinning disc confocal imaging systems were used in the study, and cells were imaged at room temperature. The first system was a Nikon Eclipse Ti-E microscope with a Yokogawa, CSU-WI spinning disc system, a Nikon LU-N4 laser launch and two Photometrics Prime BSI sCMOS cameras. The second system was a Yokogawa CSU-WI with a 100x 1.45 NA CFI Plan Apochromat Lambda objective on an Eclipse Ti2 Nikon base with a Photometrics Prime BSI sCMOS camera (as described above). For Figure 2C, cells were imaged every 5 minutes using 0.4 μm z-stack step size, and deconvolved using Nikon Elements software. For time lapse imaging in Figure 3B, cells were imaged on EMM4s agar pads and middle-focal plane images were acquired every 5 minutes. For Figure 4C, cells were mounted on glass slides and single focal planes were imaged every 200ms using continuous acquisition. The delay between frames was 657 milliseconds.

### Image analysis and quantification

To count cytoplasmic nodes, z series at 0.2 μm intervals were taken through the entire cell. Cytoplasmic clusters were counted if present within the middle 5 focal planes to avoid any cortical clusters. The number of clusters was compared to wild type through a one-way ANOVA with Tukey’s multiple comparison test (Fig S2J Mid1 graph) or a Welch’s t-test (Fig S2J Pom1 graph). Cytoplasmic fluorescence intensity was measured for Figure S2I by measuring the mean intensity in a set ROI after background subtracting. Fluorescence intensity of each strain was then compared to wild type through a one-way ANOVA with Tukey’s multiple comparison analysis. Timing of cytokinesis in Figure S3L was monitored using Sad1-mEGFP and Rlc1-mNG as respective markers of the spindle pole body and cytokinetic ring. Cells were imaged at room temperature on Mat Tek dish with YE4S agar. 0.6 μm slices through the whole cell were imaged every 3 minutes.

To quantify Cdr2 node intensity (Fig S2M) cells were grown in EMM4S at 25°and imaged under glass cover slips. 0.3 μm z-sections were taken through the whole cell and sum projections for the top half of the cell were used for quantification. The ImageJ plugin comdetv5.5 was used to identify spots. An ROI was used and spots were set to a pixel size of 4 with no intensity threshold. Large particles were set to be included and segmented. Integrated density per node is graphed for 25 cells. To quantify spots on the sides and tips of the cell (Fig S2L), single focal plane images were taken. As above, comdetv5.5 was used to identify spots. Pixel size was set at 4 and intensity was set to 10. 25 cells were analyzed. Welch’s unpaired t-test was used to analyze significance of both node intensity and node number for these experiments.

For colocalization (Fig S2A), the comdetv5.5 was used as above to identify spots from a max projection of the top half of the cell (n=25). The distance between spots was set to 3 pixels, pixel size was set as 3 and intensity was threshold at 3 for both channels. Large particles were set to be included and segmented.

To quantify Wee1 localization at nodes (Fig S3D), middle focal plane images were taken of cells grown in EMM4S at 25°. Spots were counted as nodes when associated with the membrane flanking the nucleus of interphase cells. n=100 cells for each strain performed in biological triplicate experiments. A Welch’s unpaired t-test was performed to compare wild type and *arf6Δ* cells.

For quantification of cytokinesis defects (Fig 5E), cells were stained with blankophor to highlight the division septum. Phenotypes were marked as normal, mult-septated, asymmetric, and “other,” which included tilted septa, cell wall deposits, and cells that failed to separate but initiated growth at the division site. n>80 cells for each strain performed in biological triplicate experiments.

To quantify Mid1 localization (Fig S3J), cells were marked to have nodes if spots localized to the membrane flanking the nucleus in interphase cells; n>50 cells analyzed in biological triplicate experiments. Cells were marked as abnormal (Fig S3I) if septa appeared mispositioned or contained more than the normal number of nuclei. For Cdr2 localization (Fig S3K), medial Cdr2 nodes were counted if they were found at the plasma membrane around the nucleus. n>50 cells analyzed in biological triplicate experiments. Unpaired Welch’s t-test was used for both analyses.

Cdr2 duration was quantified using a time course of single middle focal plane images in wild type or *arf6Δ* cells. Multistackreg was used to correct for cell drift. An ROI was drawn around the entire cell and Pearson’s correlation coefficient (PCC, Colocalization Plugin, FIJI) was used to compare Cdr2 localization in each frame to time point 0. A decrease in PCC over time indicates changes in the localization pattern of nodes. 10 cells were analyzed for each strain. Mean PCC and standard deviation was plotted, and Welch’s unpaired t-test was used to compare wild type and *arf6Δ* within each individual timepoint. Similar data were obtained with or without background subtraction.

### Airyscan Imaging and Analysis

An LSM880 laser scanning confocal microscope (ZEISS) with a 100x Plan Apochromat 1.46 NA oil objective, an Airyscan superresolution module, GaAsP detectors and Zen Blue acquisition software (ZEISS) was used to image Arf6 nodes (Fig 2A) and colocalization of Cdr2 and Arf6 (Fig 2B). For best resolution, 0.17 μm stacks were taken through the entire cell in resolution versus sensitivity mode. Airyscan images were then processed using the Zen Blue software.

### FRAP

FRAP analysis was performed using the LSM880 laser scanning confocal microscope described above and following previously described methods (Miller et al., 2021). Cells were imaged at a single focal plane in EMM4S (Fig 3C, S3B) or YE4S (Fig S3C) at room temperature on lectin coated μ-slide 18 well Ibidi chambers (Cat 81811). Cells were bleached by drawing an ROI within the node-containing region on one side of the cell as seen in a single middle focal plane. Two prebleach images were taken, and ROIs were bleached to <50% of the original fluorescence intensity. Unbleached cells were used to correct for photobleaching, and background was subtracted from each ROI. The intensity data were fitted using the exponential equation y = m1 + m2 * exp(-m3 * X), where m3 is the off-rate, using Prism 8 (GraphPad Software). The half-time of recovery was calculated using the equation t1/2 =(ln2)/m3. For experiments in Fig S3C, both *arf6+* and *arf6(Q75L)* were integrated into the *leu1+* locus of *arf6Δ* cells and expressed by the endogenous promoter.

### Statistical analysis

One-way ANOVA followed by Tukey’s multiple comparison test was used to assess differences for Figures 1F-G, S1C, S2I, and S2J (left panel). This test was selected to compare every mean within an experiment to each other. Unpaired t-tests with Welch’s correction were performed for Figures S1E, S2L-M, S3D, S3I-K, and for each time point in Figures 3C, S3A, and S3C. This test was selected to compare two datasets that are not assumed to have equal variance. All statistics and graphs were made using Prism8 (GraphPad Software).

### Online supplemental material

Fig. S1 relates to Fig. 1 and shows data for Arf6 controlling cell size through the Cdr2 pathway. Fig. S2 shows data for interdependent node localization of Arf6 and Cdr2. Fig. S3 contains characterization and quantification of node defects in *arf6Δ* cells. Table S1 contains yeast strains used in our study and scores from SGA screens.

## ACKNOWLEDGEMENTS

We thank members of the Moseley laboratory for discussions and comments on the manuscript; the Biomolecular Targeting Core (BioMT) (P20-GM113132) and the Imaging Facility at Dartmouth for use of equipment; and A. Paoletti for sharing strains. This project was initiated in the laboratory of Paul Nurse, whom we thank for support and advice. This work was supported by grants from the National Institute of General Medical Sciences (R01GM099774 and R01GM133856) to J.B.M.

The authors declare no competing financial interests.

